# Assessment of menstrual hygiene management and its determinants among adolescent girls: A cross-sectional study in school adolescent girls in Addis Ababa, Ethiopia

**DOI:** 10.1101/450007

**Authors:** Ephrem Biruk, Worku Tefera, Nardos Tadesse, Ashagre Sisay

## Abstract

**Introduction:** Managing menstruation is essentially dealing with menstrual flow and also in continuing regular activities like going to school, working etc. However, menstruation can place significant obstacles in girls’ access to health, education and future prospects if they are not equipped for effective menstrual hygiene management.

**Objective:** To assess the menstrual hygiene management and its determinant among school girls in Addis Ababa, Ethiopia.

**Methods:** Cross-sectional study design with quantitative method was carried out among 770 systematically selected adolescent school girls of Addis Ababa from April 1 to May 5, 2017. A self-administered pre-test close ended Amharic questionnaire at school setting was used for data collection. The coding was done using the original English version and entered to EPI-7 software. The quantitative file exported to statistical package for social science (SPSS) version 25.0 software for analysis. Total mean score was used to categorize individuals as good and poor while AOR; 95% CI with p < 0.05 was used to determine factors of menstrual hygiene management practice.

**Result:** This study had 98% response rate. 530 (70.1%) and 388(51.3%) respondents had good knowledge and practice of menstrual hygiene respectively. The findings also showed a significant positive association between good knowledge of menstruation and girls from mother’s whose education were secondary (AOR = 10.012, 95 % CI = 3.628-27.629). Wealth index quantile five (AOR = 9.038, 95 % CI = 3.728-21.909) revealed significant positive association with good practice of menstrual hygiene.

**Conclusion and recommendation:** Majority of participants had good knowledge and practice of menstrual hygiene and majority of them were from private school. Although knowledge was better than practice, girls should be educated about the process, use of proper pads or absorbents and its proper disposal.

## Introduction

### Background

Menarche is an important milestone in a girl’s transition to womanhood (1). Around the world women have developed their own personal strategies to cope with menstruation, which vary from country to country and depend on economic status, the individual’s personal preferences, local traditions and cultural beliefs and education status. The onset of menstruation presents multiple challenges for school girls. Many girls lack the knowledge, support and resources to manage menstruation in school (2).

Managing menstruation is essentially dealing with menstrual flow and also in continuing regular activities like going to school, working etc (3). However, menstruation can place significant obstacles in the way of girls’ access to health, education and future prospects if they are not equipped for effective menstrual hygiene management (MHM). Good MHM requires access to necessary resources (menstrual materials to absorb or collect menstrual blood, soap and water), facilities (private place to wash, change and dry re-usable menstrual materials, in addition to an adequate disposal system for menstrual materials), and education about MHM (1).

Schools, particularly those in developing countries, often completely lack drinking-water and sanitation and hand washing facilities; even, where such facilities exist they are often inadequate in both quality and quantity. Girls are likely to be affected in different ways from inadequate water, sanitation and hygiene conditions in schools, because the lack of such facilities they cannot attend school during menstruation (4). Girls are particularly vulnerable to dropping out of school, partly because when toilet and washing facilities are not private, not safe or simply not available in schools. Girls who reached puberty and female school staff need gender-related privacy; otherwise they may not use the facilities. This may result in absenteeism rates that can reach 10–20 per cent of school time (5).

Menstrual hygiene and management has not received adequate attention in the health and water, sanitation and hygiene (WASH) sectors in developing countries including Ethiopia and its relationship with and impact on achieving agenda of the sustainable development goal that access to adequate and equitable sanitation and hygiene for all and end open defecation, paying special attention to the needs of women and girls and those in Vulnerable situations (6).

Public investment in institutional sanitation, especially in schools and health facilities in urban areas is limited. The available sanitation facilities in most secondary schools, are poor in construction design; not convenient for sick, disabled, elderly and MHM. This in turn, results in significant unwanted impacts on health, economic activity and education (7).

According to the WHO/UNICEF Joint Monitoring Programme (JMP, 2014), the estimated coverage of urban sanitation indicated as improved, shared and other unimproved facilities have reached 27%, 40% and 26% respectively in 2015 compared to 20%, 30% and 12% respectively in 1990 (7). As a matter of fact, the existing sanitation condition for many of the school in Ethiopia is horrendous. Most school latrines are filthy and unclean, and the poor condition is contributing to high level disease prevalence, creates poor learning environments and especially impacting on girls’ education (8).

However, much attention is not given to this problem and studies on menstruation and its hygienic management as well as its influence on girls’ education are limited in Ethiopia. This study therefore, will be conducted with the aim of assessing menstrual hygiene management among primary and secondary school girls at both private and public schools.

## Objectives

### General objective

To assess the menstrual hygiene management and its determinant among private and public adolescent school girls in Addis Ababa, Ethiopia.

### Specific objective

To measure level of knowledge about menstrual hygiene management among adolescent school girls.

To measure level of menstrual hygiene management practice among adolescent school girls.

To determine factors affecting menstrual hygiene management knowledge and practice among adolescent school girls.

## Methods

### Study area

The study was conducted among primary and secondary school girls in Addis Ababa a capital of Ethiopia, from April 1 to May 15, 2017.

Addis Ababa was established in 1886 by Emperor Taitu and Minilik II and is the capital city of Federal Democratic Republic of Ethiopia. Addis Ababa is located in an area of 540.1 square kilometers and located at 9°2‘0’ North and 38°42‘0’ East at range of 2200 – 2800 meters above sea level. Despite its proximity to the equator, its lofty altitude - the third-highest capital in the world - means that it enjoys a mild climate with an average temperature of 16°C (61°F). The hottest, driest months are usually April and May. The city has administrative structures: one city council, 10 sub-cities and 116 woredas. The total population of the city projected for the year 2016, by Population Census Commission, was 3.3 million with male to female ratio of 0.92 (29).

The city has a total of 203 government and 612 private schools. Of the total number of 107,106 students enrolled from grade 7^th^ to 10^th^ education in the year 2016/17 were females (26).

### Study design

School based cross-sectional study with a quantitative research methods was employed. The survey was conducted among female adolescent students. The interviews were explored female students’ views about menstruation and its hygienic management and availability, accessibility and adequacy improved sanitation and hygiene keeping facilities for menstrual hygiene management.

## Population

### Source population

The source population of the study was all grade 7th to 10th students in public and private schools of the selected sub cities of Addis Ababa.

### Study population

The study population was all grade 7^th^ to 10^th^ students from the selected public and private schools.

### Study subjects

The study subjects were 770 randomly selected students and from whom data were collected.

## Inclusion and Exclusion criteria

### Inclusion criteria

All students who were from grade 7^th^ to 10^th^ and has monarchy was included in the study.

### Exclusion criteria

All female students who had sight problems and with mental disorders were not be included in this study.

## Sampling

### Sample size determination for quantitative study

To determine the number of adolescent school girls to be included in the study, a two-population proportion formula were used. Since the specific objectives are three, were calculated a sample size for each in order to take a large sample size.

#### Specific objective 1

The sample size of this study was determined using a single proportion formula 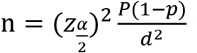 where 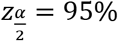 level of confidence (1.96), p= proportion of menstrual hygiene management practice among adolescent school girls in previous study (prevalence of use of sanitary napkins 35.38%) (13) and d= margin of error (5%), based on these assumption that sample size found to be 350 and to maximize the response rate of the study sample size population correction were made by multiplying design effect 2 and 10% non-response rate. Based on the above assumptions the total sample size for objective one “n” was 770 school girls.

#### Specific objective two and three

For the second objectives: factors affecting menstrual hygiene management knowledge and practice among adolescent school girls, sample size was calculated using two population proportion formula (13).

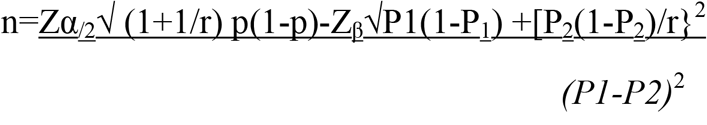

Where

Z_α/2_: 95% confidence level

Z _β_ power

P1: the probability of event in the unexposed

P2: the probability of event in the exposed

r: ratio of exposed to unexposed

OR: 1.5

It was calculated using statcalc sample size and power calculation for descriptive study of Epi info version 7

So, decision was made based on the comparison between the first specific objective (770) and second objectives (434). Finally, due to the issue of representativeness a sample size of 770 were used in the study. By using sample proportional to size, determine sample size to each school. Twenty primary and secondary schools at private and public were selected systematically from the list of schools from the selected sub-cities of Addis Ababa. A total of 770 school girls were randomly selected from students networking list of selected schools based on the proportion to the size of grade seven to ten of each school.

### Sampling procedures

The target participants for this study was adolescent school girls from grade 7^th^ to 10^th^ in very selected schools of five sub cities of Addis Ababa. A multi-stage probability sample procedure was used to select participant schools. twenty Primary and secondary schools were selected randomly from the list of schools which have grade seven to ten in the Regional Education bureau. A total of 770 School girls were randomly selected from students networking list of selected schools based on the proportion to the size of grade seven to ten of each school. The reason for the choice of school girls in grade seven to ten was because they start their menarche.

### Data collection tool and procedure

A self-administered pre-test close ended Amharic questionnaire at school setting were used. The questionnaire was contained variables related to socio-demographic characteristics, knowledge about menstruation and menstrual hygiene management, practice about menstrual hygiene. Study subjects were invited to take part voluntarily by explaining the purpose of the study and data were collected after obtaining verbal consent. Data was collected by ten female health professional data collectors with health background one supervisor.

Students were instructed on how to fill the questionnaire. Data quality was assured through careful design of the questionnaire. Data collectors and supervisor were received a one-day training on the purpose and procedure of data collection related to this research. During the training, special emphasis was given to establishing trust before asking questions. The training session were also pay attention to careful consideration for sensitive questions, observations where needed, and avoidance of participation bias. Data were checked for completeness and consistency after each day of data collection checking filled questionnaires by supervisors. The overall data collection process was coordinated by the principal investigator.

## Variables

### Dependent variables

Practice on menstrual hygiene management.

### Independent variables: -

Socio-demographic variables

Age Grade

School type

Religion

Parent’s education

Parent’s occupation.

Menstrual hygiene related Variables including:

School learning on menstrual hygiene.

Discussion with parents on menstrual hygiene.

Information before menarche on menstrual hygiene.

### Data analysis

All responses to the survey questionnaires were coded on pre-arranged coding sheet by the principal investigator to minimize errors. The coding will be using the original English version and were entered to EPI-7 software. The data file will export to statistical package for social studies (SPSS) version 25.0 software for analysis. Descriptive analysis including frequency, proportions, and measures of mean were done. Cross tabulations were made to calculate Crude and adjusted odds ratio. All variables with p<=0.20 in bivariate analysis were fitted in to the multiple logistic regression model to identify factors associated with menstrual hygienic practice. P value <= 0.05 were considered as a level of significance.

### Data quality management

The quality of data was assured at the maximum attainable level by using standardized adapted questionnaire and following the necessary procedures in order to get the intended results. To ensure quality of data, pre-test of data collection tools was done on primary school girls in New Era primary school by taking 5% of the total sample size. The data collectors were got orientation. Besides, the questionnaire was checked for completeness and correctness on daily basis by immediate supervisors.

### Operational definition

The students’ knowledge and practices was scored using a scoring system adapted from a past study. Students’ menstrual knowledge score was calculated out of the 12 knowledge specific questions (Table 3). Each correct response earned one point, whereas any wrong or don’t know response attracted no mark and thus the sum score of knowledge was calculated (12 points). Accordingly, the mean score of menstrual knowledge (7 ± 1.67) was used to decide the cutoffs of the rank. Good knowledge of menstruation and menstrual hygiene was given to those respondents who scored 7–12 points and Poor Knowledge of menstruation and menstrual hygiene was given to those respondents who scored 0–6 points. Students’ practice of menstrual hygiene score was calculated out of the practice specific questions (Table 4). Each correct response earned one point, whereas any wrong or don’t know response attracted no mark and thus the sum score of practice was calculated (15 points). And also, the mean score of menstrual practice (8± 3.619) was used to decide the cutoffs of the rank. Good practice of menstrual hygiene was given to those respondents who scored 8–15 points and poor knowledge of menstruation and menstrual hygiene was given to those respondents who scored 0–7 points. Each correct response earned one point, whereas any wrong or don’t know response attracted no mark (12).

**Hygienic menstrual management practice in school** – adolescent school girls using a clean menstrual management material to absorb or collect blood that can be changed in privacy as often as necessary for the duration of the menstruation period, using soap and water for washing the body as required, and having access to facilities to disposed of used menstrual management materials (4).

**Secondary school** – a high school or a senior high school which provides secondary education, between the ages of 14-19, after primary school and before higher education(26).

**Primary school** - a primary school or elementary school which provides primary education, between the ages of 6-14, after kinder-garden school and before secondary education(16).

**Access to water supply**– Sufficient water-collection points and water-use facilities are available in the school to allow convenient access to, and use of, water for drinking, personal hygiene, cleaning and laundry (4).

**Address the gender-related needs:** The number, location, orientation of school WASH facilities should take into consideration of the gender factor (gender mainstreaming) (8).

**Adequacy of safe water**- is the availability of 5 liters per person per day for all schoolchildren and staff at day schools(4).

**Appropriate designs for different age groups**: The detailed design of the facilities provided must also be young child friendly. Steps must be easy to climb. Door handles must be easy to reach. The toilet interior cannot be too dark. Squatting plates must be designed to accommodate a child’s feet rather than those of an adult (8).

**Improved sanitation** - are those more likely to ensure privacy and hygienic use /easily cleanable which is not full, do not have fecal matter in the squat (24)

**Physically separate facilities**: Physically separated facilities must be provided for girls, spaced sufficiently apart to ensure that girls do not feel embarrassed but secure when approaching and using the facilities. Separate hand-washing areas should also be provided, affording privacy for girls who may need to wash and dry menstrual cloths (8).

**Use appropriate orientation of facilities**: Specifically, the direction that the toilet entrance faces, must also take into account the perceived security and safety of girls. The orientation of the squatting plate should also take into account cultural and religious norms (8).

#### Ethical consideration

Ethical clearance was secured from School of Public Health, College of Health Sciences, Addis Ababa University Research Ethics Committee. Approval letter was obtained from Addis Ababa City Administration Education Bureau in the respective schools included in this study. School directors and directresses were briefed on the objectives of the study and permission to conduct the study was obtained from participating schools. The questions from the questionnaire prove not to affect the morale and personality of study subjects. Informed verbal consent was obtained from each study subject after explanation of the objective of the study. Confidentiality was ensured from all data collectors via using code numbers than names and keeping questionnaires locked. Data collectors also give health education and advice to the subjects during the data collection process.

## Result

### Socio-demographic characteristics of study population

A total of 756 primary and secondary school girls were participated from twenty primary and secondary schools, with response rate of 98%. Among these participants 38.1 % (288) were from private and the rest 61.9 % were from government schools. Among the total respondent 156 (20.6%), 160 (21.2%), 220 (29.1%) and, 220 (29.1%) were grade seven, eight, nine and ten respectively. The mean age of the study participants was 14.89 with SD ± 1.285 years, while their age range between 12-20 years. The mean age of menarche of the respondents was 12.84 with SD ± 0.745 years.

The study also indicated that 267 (35.3%) and 226 (29.9 %) of the respondents’ father completed secondary and higher level of education respectively. Regarding respondent’s mother occupation, 255 (33.7%) of them were house wife while 156 (20.6%) government employed. The majority 70.2% (531) of the respondents didn’t earn pocket money form their families. The quintile division showed that, wealth was almost equally distributed across the five quintiles. 20.5% of respondents were in the lowest quintile whereas 18.7% in the highest quintile (See table 2)

**Table 1.** Socio-demographic characteristics of primary and secondary school girls, April 1 to May 15, 2017 Addis Ababa, Ethiopia. (n =756)

| Characteristics of respondents |  | Frequency | Percentage |
| --- | --- | --- | --- |
| Religion | Orthodox | 528 | 69.8 |
|  | Muslim | 154 | 20.4 |
|  | Protestant | 59 | 7.8 |
|  | Catholic | 15 | 2.0 |
| Wealth using the wealth index | Quintile One | 155 | 20.5 |
|  | Quintile two | 148 | 19.6 |
|  | Quintile three | 151 | 20.0 |
|  | Quintile four | 161 | 21.3 |
|  | Quintile five | 141 | 18.7 |
| Mothers educational status | Illiterate | 206 | 27.2 |
|  | Read & write | 84 | 11.1 |
|  | Primary | 175 | 23.1 |
|  | Secondary | 163 | 21.6 |
|  | College and above | 128 | 16.9 |
| Live with | Both parents | 651 | 86.1 |
|  | only mother | 59 | 7.8 |
|  | Relatives | 24 | 3.2 |
|  | Only father | 22 | 2.9 |
| Fathers occupation | Merchant | 204 | 27.0 |
|  | Employed in private organization | 197 | 26.1 |
|  | Governmental employee | 180 | 23.8 |
|  | Driver | 94 | 12.4 |
|  | Daily laborer | 81 | 10.7 |

### Adolescent school girls’ knowledge about menstruation and its hygienic management

This study also found that 83 percent of respondents knew about menstruation before it occurred and their chief source of information was mothers (68.3%) followed by friends (41.08%), elder sister (32.48%) and school (28.18%). Out of the total respondents, 392 (50.9%) didn’t learn about menstrual hygiene in the school. Two hundred three (26.9%) of the respondents didn’t discuss about menstrual hygiene with their parents and friends.

Out of the total, 599 (79.2%) of girls knew that menstruation was a physiological process, whereas 42 (5.6 %) of the them believed that it was a curse from God. Majority of girls (69.3%) correctly responded hormone as the cause of menstruation. More than half, 64.6% of the respondents knew that uterus is the source of menstrual blood. (See table 3)

**Table 2.** School girls’ Information and knowledge grading on menstruation and its management, April 1 to May 15, 2017 Addis Ababa, Ethiopia. (n =756)

| Variables | Number | Percentage |
| --- | --- | --- |
| Normal menstruation cycle |  |  |
| Less than 25 days | 84 | 11.1 |
| 25 to 28 days | 515 | 68.1 |
| 28 to 35 days | 104 | 13.8 |
| More than 35 days | 53 | 7.0 |
| Normal regular menstrual bleeding duration |  |  |
| < 2 days | 52 | 6.9 |
| 2 to 7 days | 676 | 89.4 |
| >7 | 28 | 3.7 |
| Knew that there is foul smelling during menstruation |  |  |
| No | 142 | 18.8 |
| Yes | 614 | 81.2 |
| Knew that menstrual blood is unhygienic |  |  |
| No | 200 | 26.5 |
| Yes | 556 | 73.5 |
| Knew Pain during menstruation means that's one not sick |  |  |
| No | 254 | 33.6 |
| Yes | 502 | 66.4 |
| Knew menstruation is not harmful for a woman's body if she runs or dances during her period |  |  |
| No | 229 | 30.3 |
| Yes | 527 | 69.7 |
| Hear about menstruation before attaining menarche |  |  |
| No | 116 | 15.3 |
| Yes | 640 | 84.7 |
| Menstruation is not a lifelong process |  |  |
| No | 130 | 17.2 |
| Yes | 626 | 82.8 |
| Knew that a girl should take more nutritious diet during menstruation |  |  |
| No | 190 | 25.1 |
| Yes | 566 | 74.9 |

Based on the Knowledge summary of the respondent, 70.1% (67.1-73.5) had good knowledge about menstruation and its hygiene while 29.9 % of them had below the mean score of knowledge and categorized as poor knowledge.

### Menstrual Hygiene Practice

Out of the total respondents, 52.5 % (48.7-56.1) of the respondents had good practice on menstrual hygiene. Majority 457 (60.4%) of students used sanitary napkins and the rest 295 (39%) and 4 (0.5%) of them used homemade cloth and underwear as menstrual soak-up during their last menstrual period respectively. This study also enlisted the main reason for the non-use of sanitary napkins includes; 149(53.02%) high cost, 99(35.23%) difficulty in disposal, and 33(11.74%) lack of knowledge. Five hundred thirty-one participants change their pads during menstruation at school. Half 384 (59.3 %) of girls change their pads three and above times per day and 129(17.1%) changed their pads once in a day.

**Table 3.**
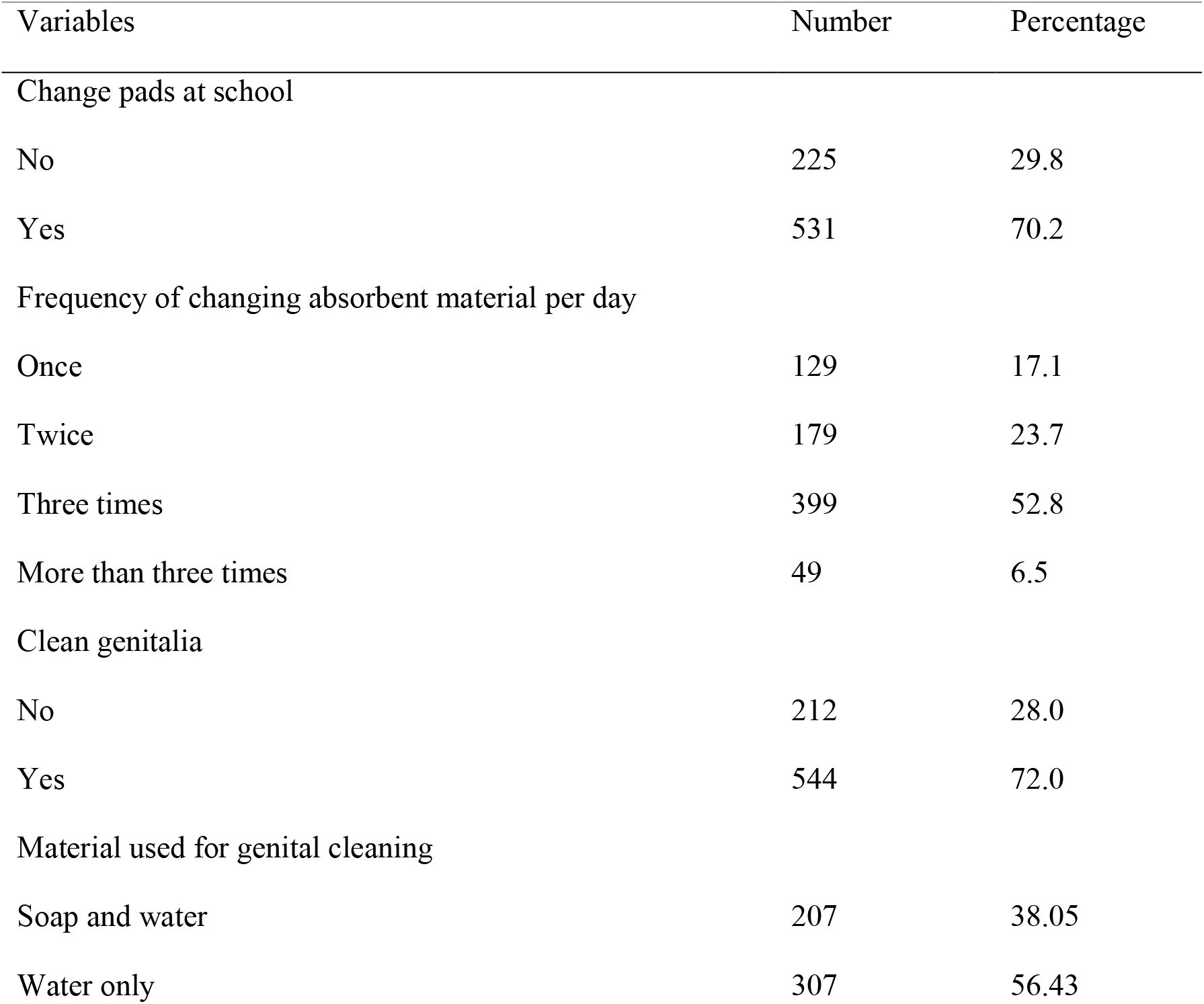
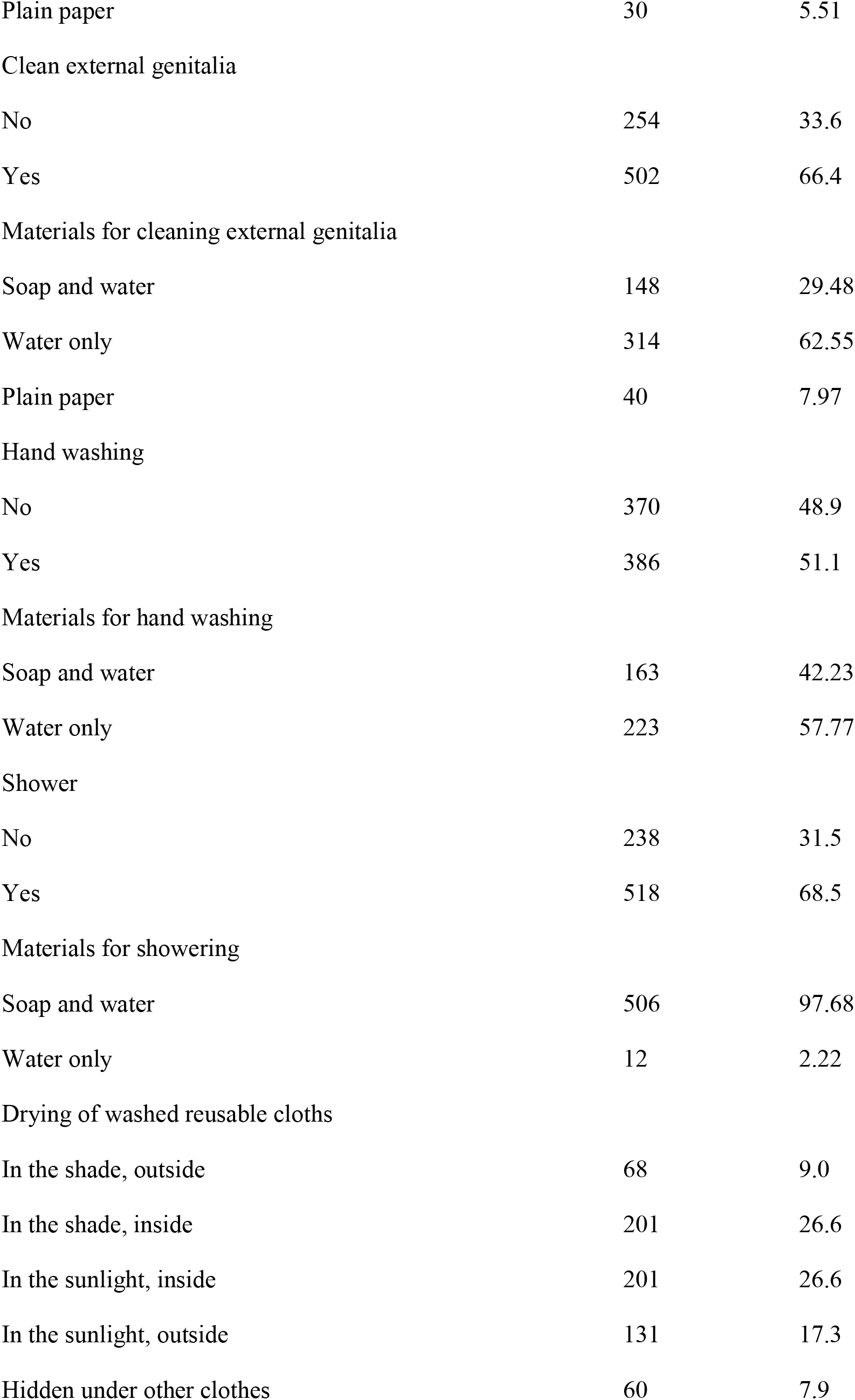
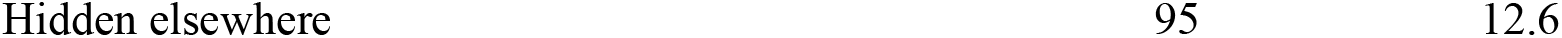
Practice grading on MHM among private and government school girl’s, April 1 to May 15, 2017 Addis Ababa, Ethiopia.

Six hundred forty (84.7 %) of the respondents disposed their used sanitary pads in the latrine, 68(9%) wrap in paper and put in the bin, and the rest 48(6.3%) threw in the open field. Three hundred thirty-one (43.9%) of the respondents dried their reusable sanitary pads in sunlight. 32.4% of the study participants store their sanitary pads in separated plastic bag for the next use. (see table 4)

This study evidenced that menstrual hygiene practice related school absenteeism was prevalent amongst respondents, 64.3% of whom miss school at least once in a month (mean 1.14, SD 1.132). Out of these respondents, 348(46.03%) of the girls were absent from school during their last menstrual period up to four days. The main reasons for school absenteeism during menstruation were; pain 294 (79.03%), followed by lack of washing facility at school 292 (78.49%), feel uncomfortable or tired 207 (55.65%), no private place to change sanitary pad 197 (48.12%), and didn’t have sanitary pad, 98 (26.34%).

Association between overall good knowledge of menstrual hygiene management and socio demographic factors

The crude value in the bivariate analysis, some of socio-demographic characteristics of the respondents were significantly associated with the outcome variable-knowledge of menstrual hygiene. The odds of good knowledge about menstruation and menstrual hygiene were 7.01 times higher for those respondents from private school compared to the government once (COR=7.01%, 95% CI: 4.936-9.952). Whereas the odds of good knowledge about menstruation and menstrual hygiene were 31.493 times higher for respondents’ mother education collage and above compared to the illiterate (COR=31.493%, 95% CI: 13.959-71.053). Unlike the crude value, the controlled model found that only the highest categories of wealth were associated with the knowledge. Girls from the wealthiest family were [AOR = 3.199, 95 % CI: 1.082-9.454] more likely to have good knowledge about menstruation and menstrual hygiene when compared to those from non-wealthy. (see table 6)

**Table 4.**
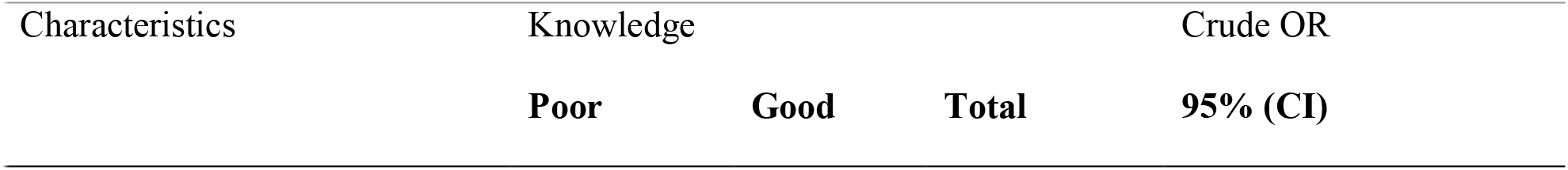
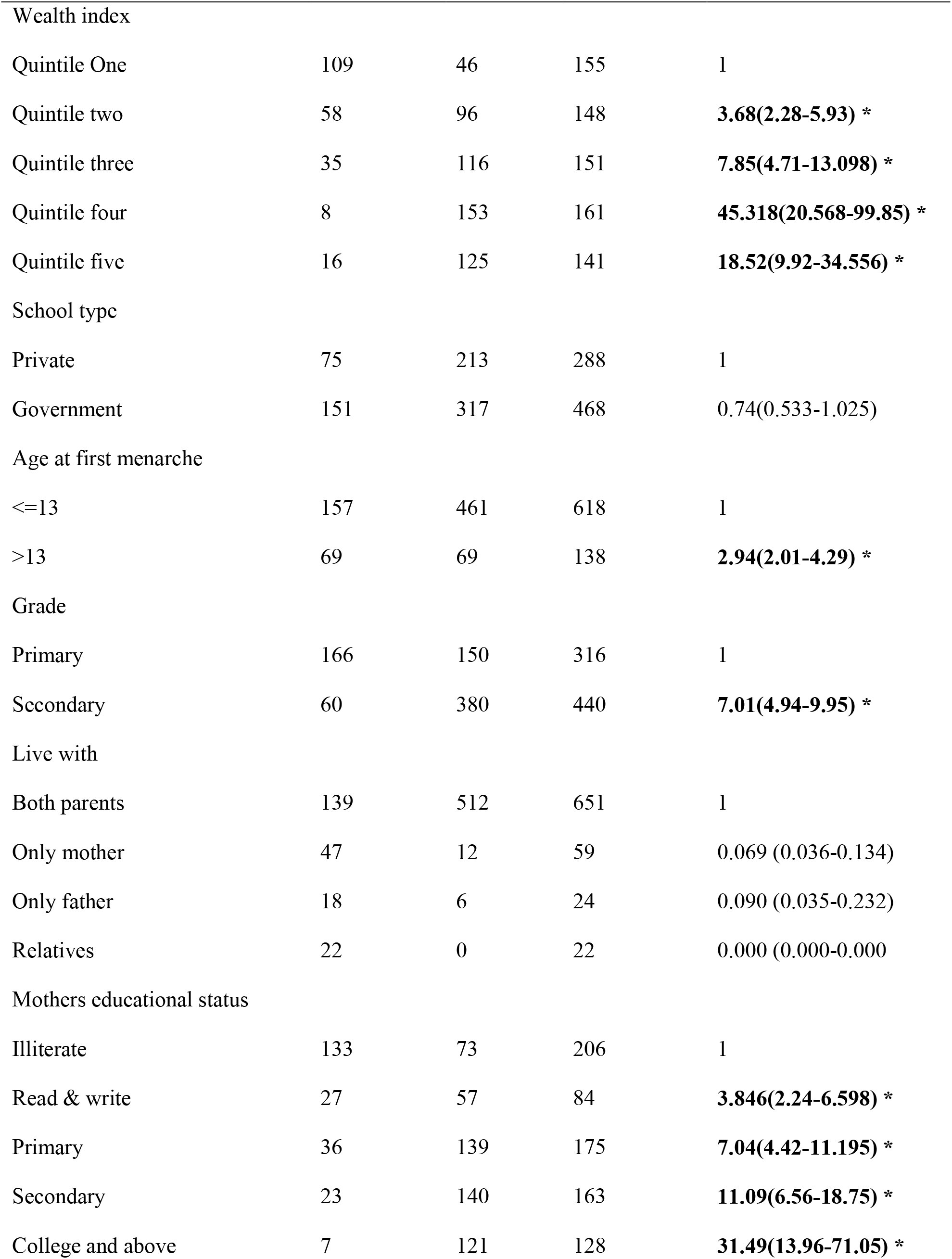
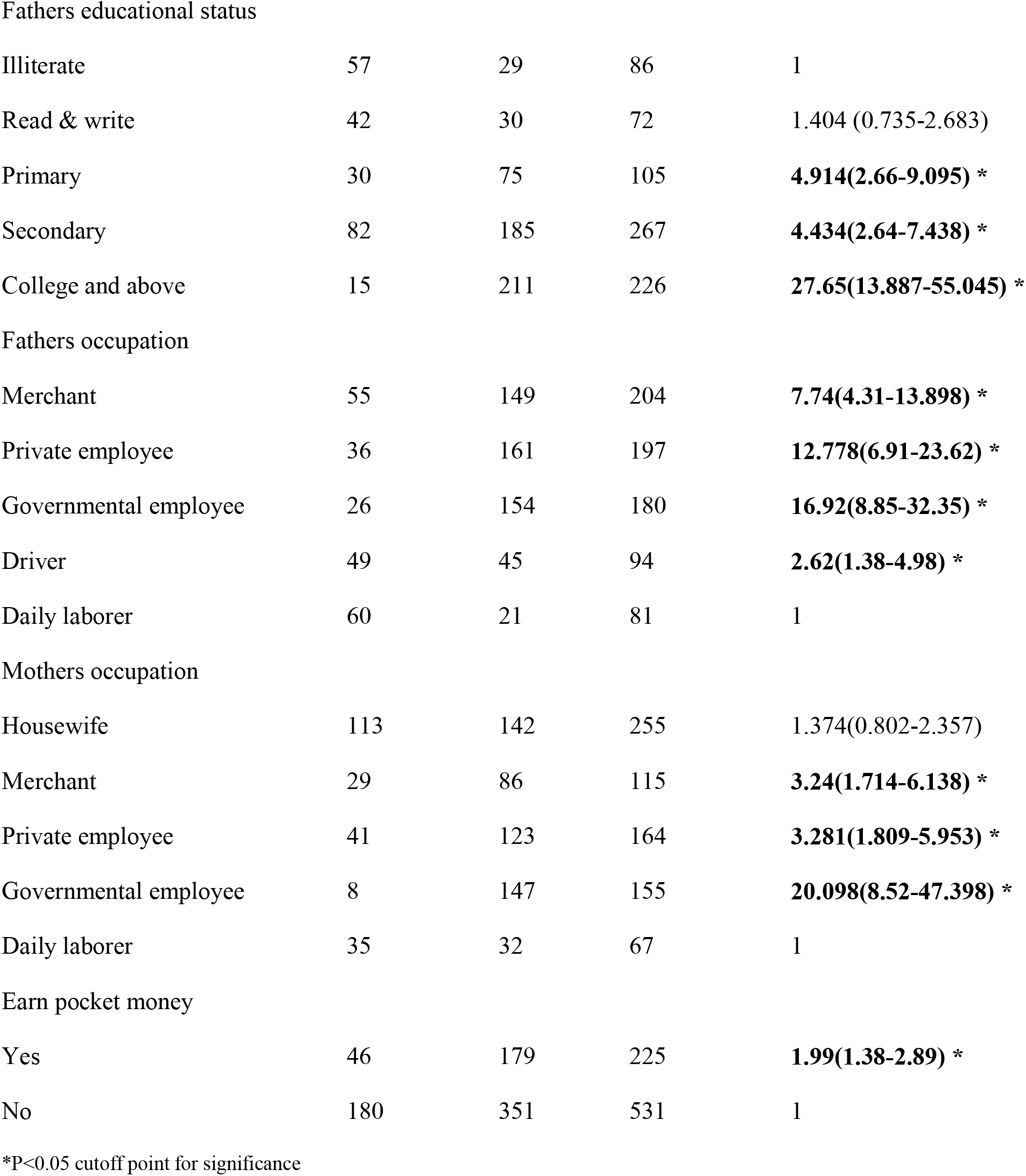
Crosstabulations between socio-demographic variables and knowledge about menstrual hygiene management among primary and secondary school girls, April 1 to May 15, 2017 Addis Ababa, Ethiopia. (n =756)

| Characteristics | Knowledge |  |  | Crude OR<br>95% (CI) |
| --- | --- | --- | --- | --- |
|  | Poor | Good | Total |  |
| Wealth index |  |  |  |  |
| Quintile One | 109 | 46 | 155 | 1 |
| Quintile two | 58 | 96 | 148 | **3.68(2.28-5.93) \*** |
| Quintile three | 35 | 116 | 151 | **7.85(4.71-13.098) \*** |
| Quintile four | 8 | 153 | 161 | **45.318(20.568-99.85) \*** |
| Quintile five | 16 | 125 | 141 | **18.52(9.92-34.556) \*** |
| School type |  |  |  |  |
| Private | 75 | 213 | 288 | 1 |
| Government | 151 | 317 | 468 | 0.74(0.533-1.025) |
| Age at first menarche |  |  |  |  |
| <=13 | 157 | 461 | 618 | 1 |
| >13 | 69 | 69 | 138 | **2.94(2.01-4.29) \*** |
| Grade |  |  |  |  |
| Primary | 166 | 150 | 316 | 1 |
| Secondary | 60 | 380 | 440 | **7.01(4.94-9.95) \*** |
| Live with |  |  |  |  |
| Both parents | 139 | 512 | 651 | 1 |
| Only mother | 47 | 12 | 59 | 0.069 (0.036-0.134) |
| Only father | 18 | 6 | 24 | 0.090 (0.035-0.232) |
| Relatives | 22 | 0 | 22 | 0.000 (0.000-0.000) |
| Mothers educational status |  |  |  |  |
| Illiterate | 133 | 73 | 206 | 1 |
| Read & write | 27 | 57 | 84 | **3.846(2.24-6.598) \*** |
| Primary | 36 | 139 | 175 | **7.04(4.42-11.195) \*** |
| Secondary | 23 | 140 | 163 | **11.09(6.56-18.75) \*** |
| College and above | 7 | 121 | 128 | **31.49(13.96-71.05) \*** |
| Fathers educational status |  |  |  |  |
| Illiterate | 57 | 29 | 86 | 1 |
| Read & write | 42 | 30 | 72 | 1.404 (0.735-2.683) |
| Primary | 30 | 75 | 105 | **4.914(2.66-9.095) \*** |
| Secondary | 82 | 185 | 267 | **4.434(2.64-7.438) \*** |
| College and above | 15 | 211 | 226 | **27.65(13.887-55.045) \*** |
| Fathers occupation |  |  |  |  |
| Merchant | 55 | 149 | 204 | **7.74(4.31-13.898) \*** |
| Private employee | 36 | 161 | 197 | **12.778(6.91-23.62) \*** |
| Governmental employee | 26 | 154 | 180 | **16.92(8.85-32.35) \*** |
| Driver | 49 | 45 | 94 | **2.62(1.38-4.98) \*** |
| Daily laborer | 60 | 21 | 81 | 1 |
| Mothers occupation |  |  |  |  |
| Housewife | 113 | 142 | 255 | 1.374(0.802-2.357) |
| Merchant | 29 | 86 | 115 | **3.24(1.714-6.138) \*** |
| Private employee | 41 | 123 | 164 | **3.281(1.809-5.953) \*** |
| Governmental employee | 8 | 147 | 155 | **20.098(8.52-47.398) \*** |
| Daily laborer | 35 | 32 | 67 | 1 |
| Earn pocket money |  |  |  |  |
| Yes | 46 | 179 | 225 | **1.99(1.38-2.89) \*** |
| No | 180 | 351 | 531 | 1 |

\*P<0.05 cutoff point for significance

The controlled effect indicated that girls whose mother’s education college and above were 6.101) [AOR, 95% CI: 1.779-20.919] times more likely to have good knowledge about menstruation and menstrual hygiene than their counterparts. The model also found that girls from secondary schools were [AOR = 13.742, 95 % CI: 5.390-35.037] more likely to have good knowledge about menstruation and menstrual hygiene when compared from primary school girls. Respondents whose mothers from government employee were strongly associated with good knowledge about menstruation and menstrual hygiene 10.555 (2.047-54.411) [AOR = 10.555, 95 % CI: 2.047-54.411]. This study also found that girls from the wealthiest family were [AOR = 3.199, 95 % CI: 1.082-9.454] more likely to have good knowledge about menstruation and menstrual hygiene when compared to those from lowest quantile. (see table 6)

**Table 5.**
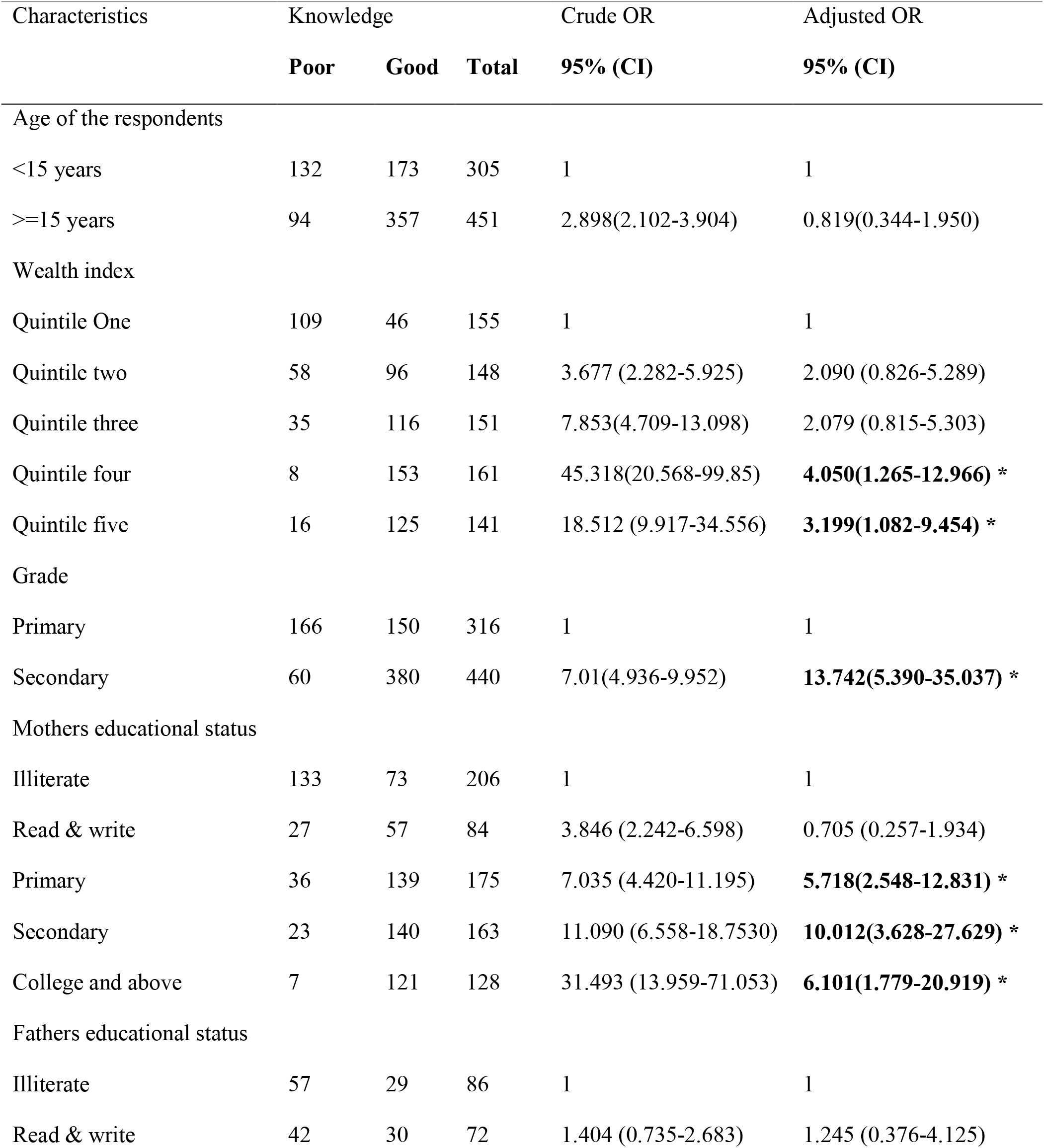
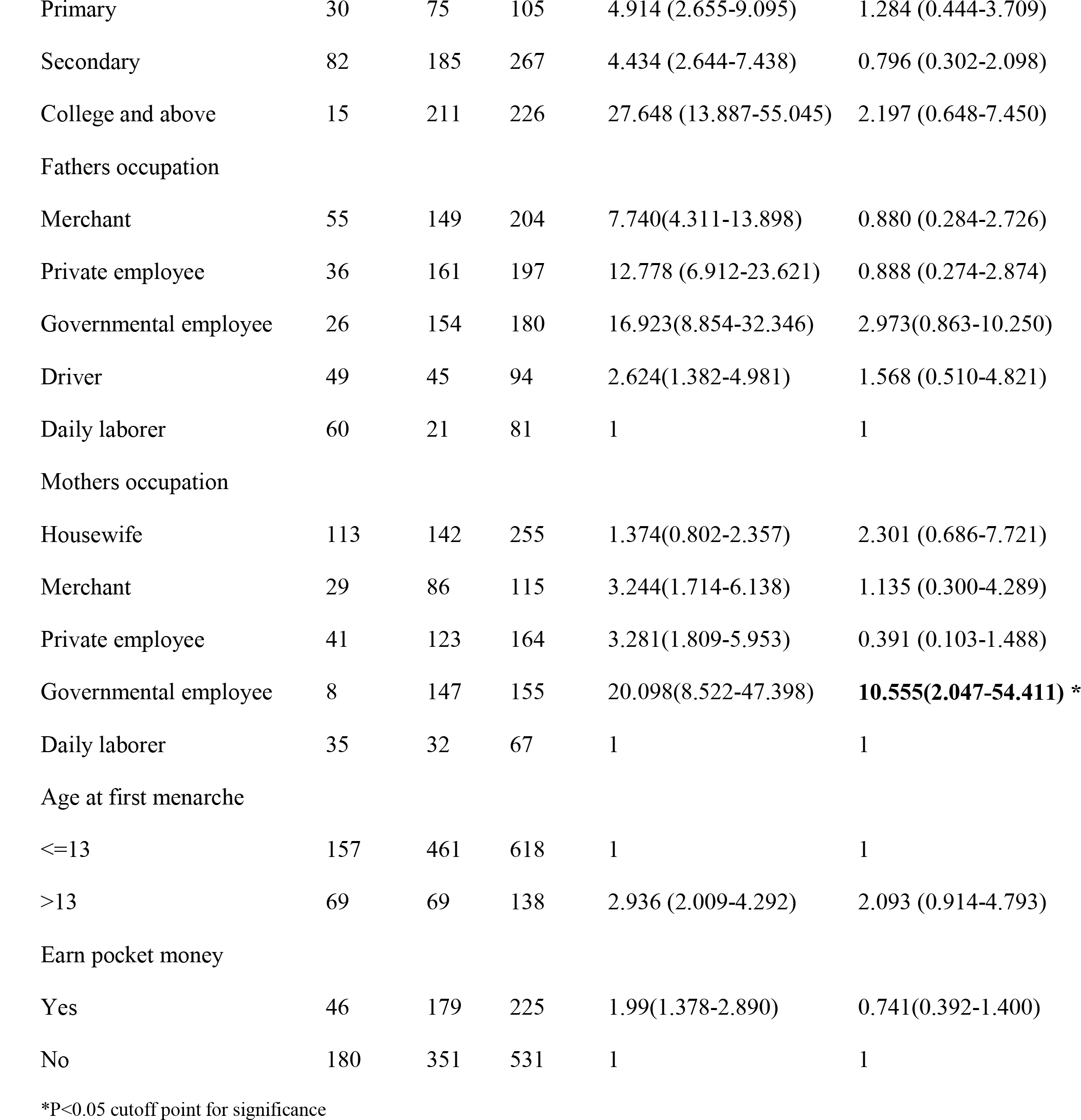
Association between socio demographic factors and knowledge about MHM among primary and secondary school girls, April 1 to May 15, 2017 Addis Ababa, Ethiopia. (n =756)

| Characteristics | Knowledge |  |  | Crude OR | Adjusted OR |
| --- | --- | --- | --- | --- | --- |
|  | Poor | Good | Total | 95% (CI) | 95% (CI) |
| Age of the respondents |  |  |  |  |  |
| <15 years | 132 | 173 | 305 | 1 | 1 |
| >=15 years | 94 | 357 | 451 | 2.898(2.102-3.904) | 0.819(0.344-1.950) |
| Wealth index |  |  |  |  |  |
| Quintile One | 109 | 46 | 155 | 1 | 1 |
| Quintile two | 58 | 96 | 148 | 3.677 (2.282-5.925) | 2.090 (0.826-5.289) |
| Quintile three | 35 | 116 | 151 | 7.853(4.709-13.098) | 2.079 (0.815-5.303) |
| Quintile four | 8 | 153 | 161 | 45.318(20.568-99.85) | **4.050(1.265-12.966) \*** |
| Quintile five | 16 | 125 | 141 | 18.512 (9.917-34.556) | **3.199(1.082-9.454) \*** |
| Grade |  |  |  |  |  |
| Primary | 166 | 150 | 316 | 1 | 1 |
| Secondary | 60 | 380 | 440 | 7.01(4.936-9.952) | **13.742(5.390-35.037) \*** |
| Mothers educational status |  |  |  |  |  |
| Illiterate | 133 | 73 | 206 | 1 | 1 |
| Read & write | 27 | 57 | 84 | 3.846 (2.242-6.598) | 0.705 (0.257-1.934) |
| Primary | 36 | 139 | 175 | 7.035 (4.420-11.195) | **5.718(2.548-12.831) \*** |
| Secondary | 23 | 140 | 163 | 11.090 (6.558-18.7530) | **10.012(3.628-27.629) \*** |
| College and above | 7 | 121 | 128 | 31.493 (13.959-71.053) | **6.101(1.779-20.919) \*** |
| Fathers educational status |  |  |  |  |  |
| Illiterate | 57 | 29 | 86 | 1 | 1 |
| Read & write | 42 | 30 | 72 | 1.404 (0.735-2.683) | 1.245 (0.376-4.125) |
| Primary | 30 | 75 | 105 | 4.914 (2.655-9.095) | 1.284 (0.444-3.709) |
| Secondary | 82 | 185 | 267 | 4.434 (2.644-7.438) | 0.796 (0.302-2.098) |
| College and above | 15 | 211 | 226 | 27.648 (13.887-55.045) | 2.197 (0.648-7.450) |
| Fathers occupation |  |  |  |  |  |
| Merchant | 55 | 149 | 204 | 7.740(4.311-13.898) | 0.880 (0.284-2.726) |
| Private employee | 36 | 161 | 197 | 12.778 (6.912-23.621) | 0.888 (0.274-2.874) |
| Governmental employee | 26 | 154 | 180 | 16.923(8.854-32.346) | 2.973(0.863-10.250) |
| Driver | 49 | 45 | 94 | 2.624(1.382-4.981) | 1.568 (0.510-4.821) |
| Daily laborer | 60 | 21 | 81 | 1 | 1 |
| Mothers occupation |  |  |  |  |  |
| Housewife | 113 | 142 | 255 | 1.374(0.802-2.357) | 2.301 (0.686-7.721) |
| Merchant | 29 | 86 | 115 | 3.244(1.714-6.138) | 1.135 (0.300-4.289) |
| Private employee | 41 | 123 | 164 | 3.281(1.809-5.953) | 0.391 (0.103-1.488) |
| Governmental employee | 8 | 147 | 155 | 20.098(8.522-47.398) | **10.555(2.047-54.411) \*** |
| Daily laborer | 35 | 32 | 67 | 1 | 1 |
| Age at first menarche |  |  |  |  |  |
| <=13 | 157 | 461 | 618 | 1 | 1 |
| >13 | 69 | 69 | 138 | 2.936 (2.009-4.292) | 2.093 (0.914-4.793) |
| Earn pocket money |  |  |  |  |  |
| Yes | 46 | 179 | 225 | 1.99(1.378-2.890) | 0.741(0.392-1.400) |
| No | 180 | 351 | 531 | 1 | 1 |

\*P<0.05 cutoff point for significance

Association between overall practice of menstrual hygiene management and socio demographic factors

Based on the bivariate analysis, among the ten socio-demographic variables, level of grade (secondary), respondent’s mother’s educational status (college and above), respondent’s father educational status (college and above), both respondent’s mother and father occupational status, Age at first menarche (>=13) and pocket money were significantly associated with 95 % CI COR at P<0.05 with overall practice of menstrual hygiene management among respondents.

The crude value also showed that the odds of overall practice of menstrual hygiene management among secondary school girls were 2.364 (95% C.I: 1.759-3.177) times higher compared to primary school girls. In this study, it was found that pocket money was associated with overall practice of menstrual hygiene management. The odds of overall practice of menstrual hygiene management among girls 2.177 (95% C.I: 1.575-3.009) times higher for respondent who had earned pocket money from their families. The odds of overall practice of menstrual hygiene management among girls 2.330 (95% C.I: 1.720-3.156) times higher among respondent who were from private school than government school girls. (see table 7)

**Table 6.** Crosstabulations between socio-demographic variables and MHM practice among primary and secondary school girls, April 1 to May 15, 2017 Addis Ababa, Ethiopia. (n =756)

| Characteristics | Practice |  |  | Crude OR |
| --- | --- | --- | --- | --- |
|  | Poor | Good | Total | 95% (CI) |
| School type |  |  |  |  |
| Government | 259 | 209 | 468 | 1 |
| Private | 100 | 188 | 288 | **2.33(1.72-3.16) \*** |
| Age of the respondents |  |  |  |  |
| <15 years | 180 | 125 | 305 | 1 |
| >=15 years | 179 | 272 | 451 | **2.19(1.63-2.94) \*** |
| Wealth index |  |  |  |  |
| Quintile One | 133 | 22 | 152 | 1 |
| Quintile two | 91 | 57 | 148 | **3.79(2.16-6.63) \*** |
| Quintile three | 74 | 77 | 151 | **6.29(3.620-10.931) \*** |
| Quintile four | 32 | 129 | 161 | **24.37(13.45-44.16) \*** |
| Quintile five | 29 | 112 | 164 | **23.35(12.71-42.91) \*** |
| Grade |  |  |  |  |
| Primary | 189 | 127 | 316 | 1 |
| Secondary | 170 | 270 | 440 | **2.36(1.76-3.18) \*** |
| Religion |  |  |  |  |
| Orthodox | 282 | 246 | 528 | 1 |
| Muslim | 40 | 114 | 154 | 0.582(0.204-1.657) |
| Protestant | 31 | 28 | 59 | 1.900(0.636-5.674) |
| Catholic | 6 | 9 | 15 | 0.602(0.19-1.906) |
| Live with |  |  |  |  |
| Both parents | 269 | 382 | 651 | 1 |
| Only mother | 44 | 15 | 59 | 0.280(0.131-0.440) |
| Only father | 24 | 0 | 24 | 0.998(0.000-0.000) |
| Relatives | 22 | 0 | 22 | 0.998(0.000-0.000) |
| Mothers' educational status |  |  |  |  |
| Illiterate | 150 | 56 | 206 | 1 |
| Read & write | 42 | 42 | 84 | **2.68(1.58-4.54) \*** |
| Primary | 100 | 75 | 175 | **2.01(1.31-3.08) \*** |
| Secondary | 48 | 115 | 163 | **6.42(4.07-10.12) \*** |
| College and above | 19 | 109 | 128 | **15.37(8.64-27.35) \*** |
| Fathers' educational status |  |  |  |  |
| Illiterate | 64 | 22 | 86 | 1 |
| Read & write | 49 | 23 | 72 | 1.365(0.683-2.730) |
| Primary | 39 | 66 | 105 | **4.92(2.634-9.203) \*** |
| Secondary | 135 | 132 | 267 | **2.84(1.657-4.884) \*** |
| College and above | 72 | 154 | 226 | **6.22(3.556-10.887) \*** |
| Fathers' occupation |  |  |  |  |
| Merchant | 74 | 130 | 204 | **9.19(4.757-17.750) \*** |
| Private employee | 55 | 142 | 197 | **13.51(6.911-26.391) \*** |
| Governmental employee | 86 | 94 | 180 | **5.717(2.951-11.078) \*** |
| Driver | 76 | 18 | 94 | 1.239(0.565-2.716) |
| Daily laborer | 68 | 13 | 81 | 1 |
| Mothers' occupation |  |  |  |  |
| Housewife | 145 | 110 | 255 | **2.63(1.407-4.92) \*** |
| Merchant | 41 | 74 | 115 | **6.26(3.140-12.47) \*** |
| Private employee | 63 | 101 | 164 | **5.56(2.887-10.699) \*** |
| Governmental employee | 58 | 97 | 155 | **5.798(2.996-11.22) \*** |
| Daily laborer | 52 | 15 | 67 | 1 |
| Age at first menarche |  |  |  |  |
| <=13 | 268 | 350 | 618 | 1 |
| >13 | 91 | 47 | 138 | **2.53(1.718-3.72) \*** |
| Earn pocket money |  |  |  |  |
| Yes | 77 | 148 | 225 | **2.18(1.575-3.01) \*** |
| No | 282 | 249 | 531 | 1 |

\*P<0.05 cutoff point for significance.

After controlling interaction effect of all the variables, it was found that the current age of school girl students above fifteen years old were 2.283 [AOR (95% C.I: 1.613-4.971)] times more likely to have good practice than their counterparts. Girls whose mother’s education secondary and above were eight times more likely to have good practice about menstrual hygiene compared to those from illiterate mothers [AOR = 7.761, 95 % CI: 3.583-16.809)]. Girls whose fathers from private employee were [AOR (95% C.I): 3.654 (1.215-10.991) times more likely had good menstrual hygiene management practice than those who were daily laborer family. This study also found that girls whose age at first menarche greater than thirteen were 2.572 times more likely to have good practice about menstrual hygiene compared to those who were less than thirteen years old. [AOR (95% C.I): 2.572 (1.409-4.694)]. Practice of MHM was also distributed unequally across the wealth quintile. The odd of good practices were 3 times higher in the highest quintile compared to the 2^nd^. In addition, the overall knowledge of the respondents was significantly associated with their practice [AOR (95% C.I): 4.581 (2.462-8.526)] (See table 8)

**Table 7.** Association between socio demographic variables and level of MHM practice among primary and secondary school girls, April 1 to May 15, 2017 Addis Ababa, Ethiopia. (n =756)

| Characteristics | Practice |  |  | Crude OR | Adjusted OR |
| --- | --- | --- | --- | --- | --- |
|  | Poor | Good | Total | 95% (CI) | 95% (CI) |
| School type |  |  |  |  |  |
| Government | 259 | 209 | 468 | 1 | 1 |
| Private | 100 | 188 | 288 | 2.330(1.720-3.156) | 1.09(0.64-1.85) |
| Age of the respondents |  |  |  |  |  |
| <15 years | 180 | 125 | 305 | 1 | 1 |
| >=15 years | 179 | 272 | 451 | 2.188(1.627-2.942) | **2.832(1.62-4.97) \*** |
| Wealth index |  |  |  |  |  |
| Quintile One | 133 | 22 | 152 | 1 | 1 |
| Quintile two | 91 | 57 | 148 | 3.787 (2.164-6.626) | **2.968(1.32-6.67) \*** |
| Quintile three | 74 | 77 | 151 | 6.291 (3.620-10.931) | **3.80(1.730-8.35) \*** |
| Quintile four | 32 | 129 | 161 | 24.371 (13.45-44.16) | **14.776(6.23-35.04) \*** |
| Quintile five | 29 | 112 | 164 | 23.348 (12.71-42.91) | **9.038(3.728-21.91) \*** |
| Grade |  |  |  |  |  |
| Primary | 189 | 127 | 316 | 1 | 1 |
| Secondary | 170 | 270 | 440 | 2.364(1.759-3.177) | **3.115 (1.606-6.04) \*** |
| Mothers' educational status |  |  |  |  |  |
| Illiterate | 150 | 56 | 206 | 1 | 1 |
| Read & write | 42 | 42 | 84 | 2.679(1.582-4.535) | 1.588(0.71-3.560) |
| Primary | 100 | 75 | 175 | 2.009(1.308-3.084) | 1.929(0.95-3.933) |
| Secondary | 48 | 115 | 163 | 6.417(4.069-10.122) | **7.761(3.58-16.81) \*** |
| College and above | 19 | 109 | 128 | 15.367(8.639-27.352) | **17.91(6.65-48.22) \*** |
| Fathers' educational status |  |  |  |  |  |
| Illiterate | 64 | 22 | 86 | 1 | 1 |
| Read & write | 49 | 23 | 72 | 1.365(0.683-2.730) | 0.482(0.146-1.588) |
| Primary | 39 | 66 | 105 | 4.923(2.634-9.203) | 0.552(0.218-1.403) |
| Secondary | 135 | 132 | 267 | 2.844(1.657-4.884) | 0.570(0.110-1.663) |
| College and above | 72 | 154 | 226 | 6.222(3.556-10.887) | 0.574(0.099-1.761) |
| Fathers' occupation |  |  |  |  |  |
| Merchant | 74 | 130 | 204 | 9.189(4.757-17.750) | 1.933(0.705-5.295) |
| Private employee | 55 | 142 | 197 | 13.505(6.911-26.391) | **3.65(1.22-10.991) \*** |
| Governmental employee | 86 | 94 | 180 | 5.717(2.951-11.078) | 1.178(0.413-3.363) |
| Driver | 76 | 18 | 94 | 1.239(0.565-2.716) | 0.481(0.166-1.391) |
| Daily laborer | 68 | 13 | 81 | 1 | 1 |
| Mothers' occupation |  |  |  |  |  |
| Housewife | 145 | 110 | 255 | 2.630(1.407-4.916) | 2.125 (0.759-5.952) |
| Merchant | 41 | 74 | 115 | 6.257(3.140-12.470) | 1.570 (0.522-4.718) |
| Private employee | 63 | 101 | 164 | 5.558(2.887-10.699) | 1.259 (0.448-3.539) |
| Governmental employee | 58 | 97 | 155 | 5.798(2.996-11.219) | 0.813 (0.272-2.435) |
| Daily laborer | 52 | 15 | 67 | 1 | 1 |
| Age at first menarche |  |  |  |  |  |
| <=13 | 268 | 350 | 618 | 1 | 1 |
| >13 | 91 | 47 | 138 | 2.529(1.718-3.721) | **2.57(1.41-4.69) \*** |
| Earn pocket money |  |  |  |  |  |
| Yes | 77 | 148 | 225 | 2.177(1.575-3.009) | 1.11(0.67-1.81) |
| No | 282 | 249 | 531 | 1 | 1 |

\*P<0.05 cutoff point for significance.

Association between overall knowledge and practice of menstrual hygiene management and other menstrual related factors

The regression model also evidenced other menstrual related factors like learning and discussing about MHM in the school and with parents and friend and also hearing about it before menarche were significantly associated with both the outcome variables. The odds of good knowledge about menstrual hygiene management among those who learn about menstrual hygiene at school were 3.110 (95% CI: 1.569-6.162) times higher than those who didn’t learn at their school. (See table 9)

**Table 8.** Association between other related variables and knowledge about MHM among primary and secondary school girls, April 1 to May 15, 2017 Addis Ababa, Ethiopia. (n=756)

| Characteristics | Knowledge |  | Crude OR | Adjusted OR |
| --- | --- | --- | --- | --- |
|  | Good | Poor | 95% (CI) | 95% (CI) |
| Learn about menstrual hygiene in |  |  |  |  |
| the school |  |  |  |  |
| Yes | 316 | 55 | 4.59(3.24-6.51) | **3.11(1.57-6.16) \*** |
| No | 214 | 171 | 1 | 1 |
| Discuss about menstrual hygiene with their parents |  |  |  |  |
| Yes | 481 | 72 | 20.99(13.99-31.51) | **10.89(5.39-21.98) \*** |
| No | 49 | 154 | 1 | 1 |
| Heard about menstruation before attaining menarche |  |  |  |  |
| Yes | 500 | 140 | 10.24(6.49-16.15) | **5.03(2.21-11.48) \*** |
| No | 30 | 86 | 1 | 1 |

\*P<0.05 cutoff point for significance.

This study found that the odds of good practice about menstrual hygiene management among those who discuss about menstrual hygiene with their parents were 13.651 (95% CI: 7.087-26.296) times higher than those who didn’t discuss about menstrual hygiene with their parents. (See table 10)

**Table 9.**
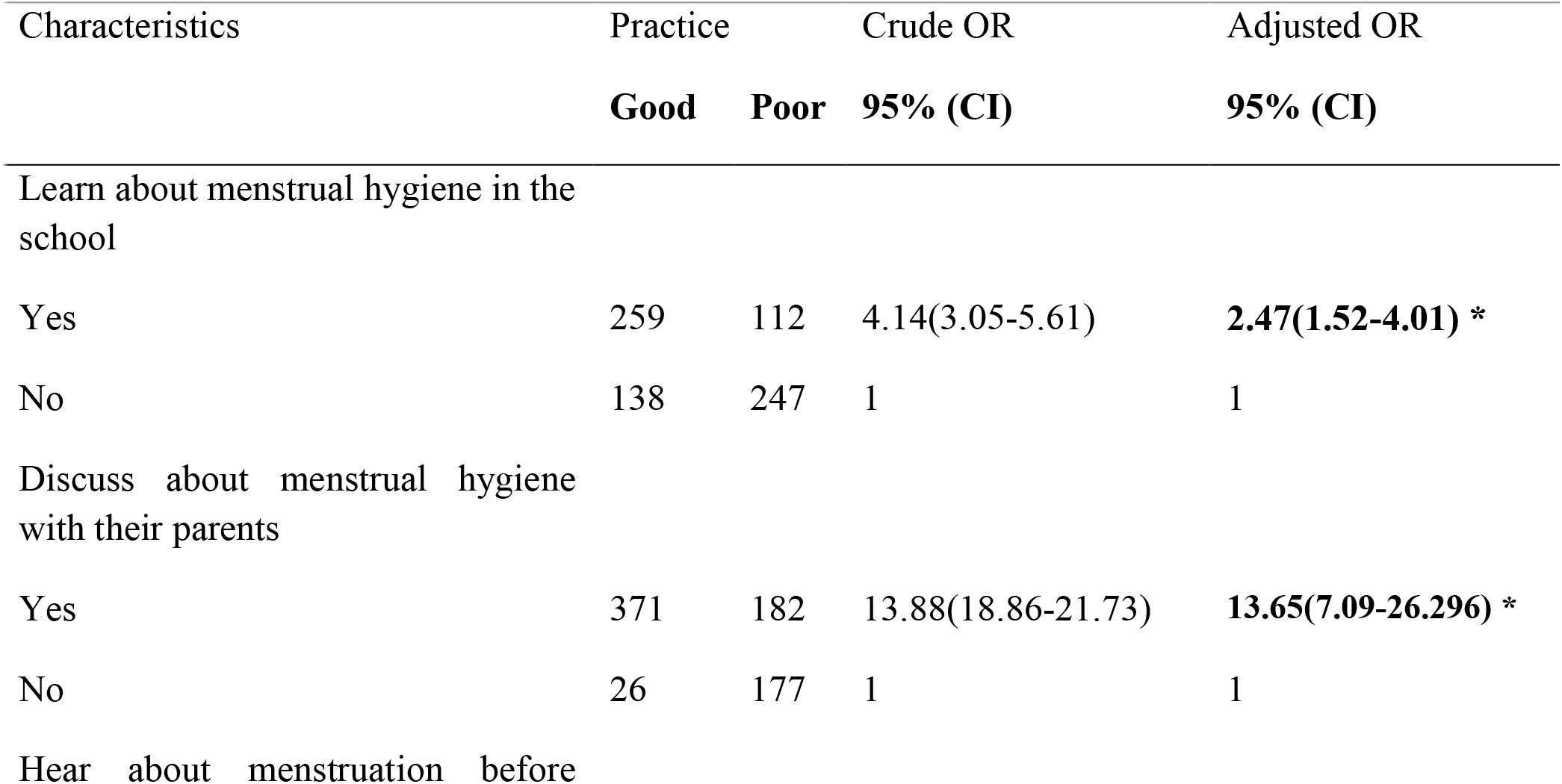
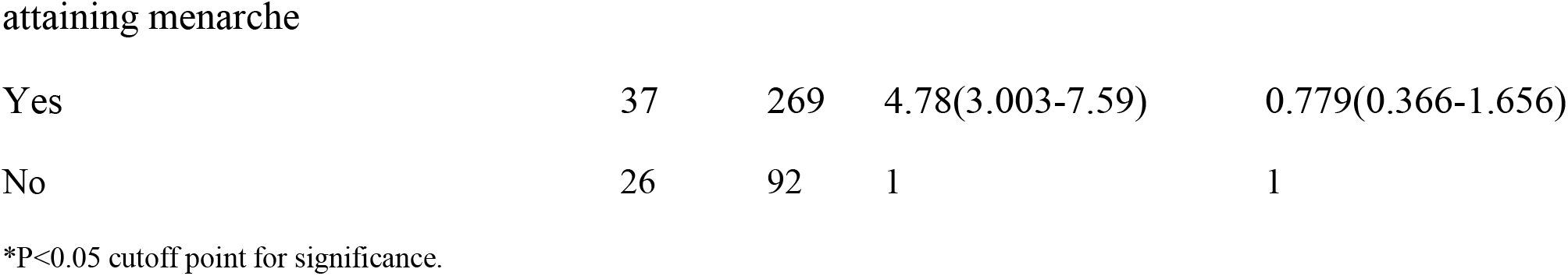
Association between socio demographic factors and MHM practice among primary and secondary school girls in Addis Ababa, 2017. (n=756)

## Discussion

The onset of menstruation is one of the most important changes occurring among the girls during the adolescent years. The bodily changes associated with puberty will have an impact in the girl’s physical, psychological and social development (19–21). This study was conducted to identify factors affecting practice of menstrual hygiene among school girls in Addis Ababa.

In this study, the mean age of menarche of the respondents was 12.84 with SD +0.745 years which is similar to studies conducted in Jammu district, India and Argoha village of Haryana with the mean age of menarche 13.43±.83 and 12.76 ± 0.936 years respectively (30,31).

In consistent with study reports from Jimma district, India (66.15%) (30), for about 68% school girls, their mothers were main source of information on menstruation. This could be suggestive of the contribution of mothers for hygienic practice of girls during menarche. In contrast to this finding, school teachers and sisters respectively, were reported to be important source of such information in Amhara North East, Ethiopia (13).

### Factors of menstrual hygiene management

Present study also reported that, majority (70.1 %) of the students had good knowledge about menstruation and menstrual hygiene. The finding was similar with the result from studies done in western Ethiopia; 60.9%. This study notified that Five hundred ninety-nine (79.2%) of girls knew that menstruation to be normal a physiological process. Such prevalence found to be consistent with a result of a study done in western Ethiopia (76.9%), Argoha village of Haryana 71.3%, and Central India 89% of school girls knew correctly that menstruation as physiologic process(12,31,32). A possible explanation for this similarity may be that girls had good discussion in families openly. This finding however, was higher than that of those in previous a study done in Northeast Ethiopia; 319 (57.89%) This difference might be due to the study method difference (mixed qualitative and quantitative method for the previous study (12).

Adolescent girls who knew that uterus was the source of blood in menstruation were 64.6% which is similar to studies conducted in Amhara Ethiopia, western Ethiopia, and Central India was found out to be 60.9%, 59.3% and 60% respectively (12,32,33). This study disagrees with results obtained from a study in Argoha village of Haryana 38.7%, possibly due to minimum information provided about menstruation and menstrual hygiene by schools and families (31).

In this study, multivariable analysis showed that girls whose mother’s educational status secondary school and above were 10.012 times more likely to had good knowledge about menstruation and menstrual hygiene than their counterparts [AOR = 10.012, 95 % CI: 3.628-27.629]. A similar study done in western Ethiopia and Jammu District India showed that, parental education was positively associated with girls’ menstrual knowledge (12,30). The reason could that educated mothers may provide information about menstruation and menstrual hygiene to their daughters. Girls from educated families may discuss openly about menstruation.

Unlike a study done in Amhara region, Ethiopia 2016 [AOR = 0.94, 95 % CI: 0.46–1.92], this study found that, the grade level of respondents was positively and significantly [AOR = 13.74, 95 % CI: 5.39-35.04] associated with knowledge about menstruation and menstrual hygiene management. Due to the fact that high school girls might have high possibility for exposure to information regarding menstruation and its hygienic management (11). in another study done in Odisha, India 2015, none of the mother’s occupational category were significantly related with knowledge (25). But in this study only government employed mothers were the one associated with knowledge on menstruation.

Regarding hygiene related practices during menstruation, this study found that 9.27% girls took daily bath during menstruation and 29.48% clean their external genital with soap and water during menstruation. A similar study done in Jammu and Kashmir, India indicated that 93.18% had daily bath and 66.67% clean external genital with soap and water (30). The difference might be due to socio cultural, weather condition and economic factors.

In this study, three hundred ninety-seven (52.5%) of the respondents had good practice of menstrual hygiene. The finding of this study was lower than studies conducted in the Amhara region of Ethiopia and Jammu and Kashmir, India which were 84.28 % and 59.09%, respectively (30,33). Comparatively, lower level of practice of menstrual hygiene was recorded from similar study conducted on high school girls in Western Ethiopia, it was indicated that only 39.9 % of the study participants practice good menstrual hygiene (12). Thus, the reason for the observed difference could be due to low awareness and communication of menstrual hygiene by high school Western Ethiopia girls which affects their menstrual hygienic practice.

In this study, it was found that 17.3% of the study participants dried their reusable pads in the sun light outside which is similar with the finding in Amhara, Ethiopia 15.5% of the participants dried in the sun light. In contrast, a study done in central India indicated that 93% of adolescent girls dried reusable pads in the sun light (32,33). This difference might be due to different in the socio-cultural factors. 457(60.4%) of participants use disposable sanitary pads during menses. In a similar study done in Argoha village of Haryana and central India Indicated that, 80.7 % and 98% girls use only napkin during menses respectively (31,32). Comparatively, lower level of use of sanitary pads was recorded from similar study conducted on high school girls in north east Ethiopia, it was indicated that 35.38% of the study participants use disposable sanitary pads during menses. This may be due to differences in socio economic differences. The main reason for not using pads in present study was non-affordability due to high cost (53.02%) followed by non-availability and disposal problems which is similar to study in Jammu district India(78.94%) (30).

The adjusted value of this study found that, both mother’s educational category was [AOR = 7.761, 95 % CI: 3.583-16.809] positively associated with practice of menstrual hygiene which agrees with the study done in western Ethiopia, in 2014 and Northeast Ethiopia [AOR = 2.03, 95 % CI: 1.38–2.97] [AOR = 4.26, 95 % CI: 1.61 - 11.28] respectively (11,12). Multivariable analysis showed that girls who learn and discuss about menstrual hygiene in their school and with their parent were 2.472, and 13.651 times more likely to had good practice about menstruation and menstrual hygiene than their counterparts. A similar study done in northeast Nigeria that learn about menstrual hygiene in the school was positively associated with girl’s menstrual practice (34). Possibly due to information provided about menstrual hygiene management at schools. Students whose age above fifteen were 2.832 times more likely to have good practice than age less than fifteen a similar study done in northeast Nigeria. A significant association was also observed between girls whose first menarche were above thirteen (AOR= 2.572) and below with the practice of MHM. Which was also significant in another study done in south India (24).

## Conclusion

Seventy percent of the participants had good knowledge of menstruation and menstrual hygiene and it was better among private school girls than the government. Half of the total respondents had good practice of menstrual hygiene among respondents, and alike to that of knowledge, the large proportion of them were from private school.

Good knowledge of menstruation showed a significantly positive association with the level of grade, educational status of the mother, and mother’s occupational status. Except for the level of grade, all the above variables plus current age of respondents’ and age at first menarche were positively associated with practice of menstrual hygiene management.

All those factors considered in addition to socio demographic variables, which assume to be predictive of the outcome variables including learn about menstrual hygiene in the school, discuss about menstrual hygiene with their parents and friends and hear about menstruation before attaining menarche were positively associated.

## Recommendation

Federal level managers are recommended to:

Strengthen the enabling environment through advocacy and policy initiatives for improved WASH and MHM education.

Promote an innovative, intercultural, multi-sectorial and gender approach in all programming, ensuring that MHM aspects are included in planning processes and budget allocation processes by the water and sanitation, health, and education ministries.

Regional Level: in addition to federal recommendations, regional level managers are advised to:

Give technical assistance and advocacy to prioritize budgets and investment in WASH facilities in schools by health, and education bureau.

Strengthen teachers’ capacities and equips them with tools to provide in-depth and medically accurate information to students in a safe learning environment by education bureau.

Strengthen school health packages provided by health extension professionals by health bureau.

School level managers suggested to:

Establish coordination between students, teachers and parents to improve MHM conditions at schools.

Complement menstrual hygiene management as part of the school health programs and should also give special attention towards making schools a comfortable place for girl’s menstrual hygiene practice by continuous provision of sanitary pad especially for the neediest ones.

Consult parents about the need to support their children with sanitary materials for menstrual hygiene in addition to other basic hygienic products during parent-school teacher meeting.

Educate and counsel girls about the important and the need for good personal hygiene including hand washing practice during menstruation by using peer group discussion which supposedly mediated by female school teachers.

Parents are advised to:

Educate their daughters about the process, good personal hygiene, use of proper pads, and its proper disposal.

Support their children with sanitary materials for menstrual hygiene.

## Acknowledgment

It gives me pleasure to express my heartfelt gratitude to Addis Ababa University, School of Public Health librarians for providing me the necessary references. I would like to thank to my advisor Worku Tefera for his unrestricted guidance, envisioning and supporting from the period of instigation to this project.

I am also grateful to thank Addis Ababa education bureau, sub city education offices, school staffs, my office staffs for their cooperation and assistance in giving important information.

I am pleased to extend my thanks to my brother, Mehary Biruk, for his devotion to support me through the academic years.

I would also like to thank all the participants of the study for their patience and collaboration during data collection.

I am exclusively thankful to my friends Ashagre Sisay, Ermias Woldie, Tasew Denbi, Nardos Tadesse and Sintayehu Tadesse for their vital contribution for success of this work.

